# Genetic conflict with a parasitic nematode disrupts the legume-rhizobia mutualism

**DOI:** 10.1101/213876

**Authors:** Corlett W. Wood, Bonnie L. Pilkington, Priya Vaidya, Caroline Biel, John R. Stinchcombe

## Abstract

Genetic variation for partner quality in mutualisms is an evolutionary paradox. One possible resolution to this puzzle is that there is a tradeoff between partner quality and other fitness-related traits. Here, we tested whether a susceptibility to parasitism is one such tradeoff in the mutualism between legumes and nitrogen-fixing bacteria (rhizobia). We performed two greenhouse experiments with the legume *Medicago truncatula*. In the first, we inoculated each plant with the rhizobia *Ensifer meliloti* and with one of 40 genotypes of the parasitic root-knot nematode *Meloidogyne hapla*. In the second experiment, we inoculated all plants with rhizobia and half of the plants with a genetically variable population of nematodes. Using the number of nematode galls as a proxy for infection severity, we found that plant genotypes differed in susceptibility to nematode infection, and nematode genotypes differed in infectivity. Second, we showed that there was a genetic correlation between the number of mutualistic structures formed by rhizobia (nodules) and the number of parasitic structures formed by nematodes (galls). Finally, we found that nematodes disrupt the rhizobia mutualism: nematode-infected plants formed fewer nodules and had less nodule biomass than uninfected plants. Our results demonstrate that there is genetic conflict between attracting rhizobia and repelling nematodes in *Medicago*. If genetic conflict with parasitism is a general feature of mutualism, it could account for the maintenance of genetic variation in partner quality and influence the evolutionary dynamics of positive species interactions.

**Impact summary:** Cooperative species interactions, known as mutualisms, are vital for organisms from plants to humans. For example, beneficial microbes in the human gut are a necessary component of digestive health. However, parasites often infect their hosts via mechanisms that are extraordinarily similar to those used by mutualists, which may create a tradeoff between attracting mutualists and resisting parasites. In this study, we investigated whether this tradeoff exists, and how parasites impact mutualism function in the barrelclover *Medicago truncatula*, a close relative of alfalfa. Legumes like *Medicago* depend on nitrogen provided by mutualistic bacteria (rhizobia) to grow, but they are also infected by parasitic worms called nematodes, which steal plant nutrients. Both microorganisms live in unique structures (nodules and galls) on plant roots. We showed that the benefits of mutualism and the costs of parasitism are predicted by the number of mutualistic structures (nodules) and the number of parasitic structures (galls), respectively. Second, we found that there is a genetic tradeoff between attracting mutualists and repelling parasites in *Medicago truncatula*: plant genotypes that formed more rhizobia nodules also formed more nematode galls. Finally, we found that nematodes disrupt the rhizobia mutualism. Nematode-infected plants formed fewer rhizobia nodules and less total nodule biomass than uninfected plants. Our research addresses an enduring evolutionary puzzle: why is there so much variation in the benefits provided by mutualists when natural selection should weed out low-quality partners? Tradeoffs between benefits provided by mutualists and their susceptibility to parasites could resolve this paradox.

## Introduction

Nearly all species require mutualists to carry out crucial biological functions (Shapira 2016). Insects partner with mutualists for nutrition (Hansen and Moran 2014; Nygaard et al. 2016); most plants rely on mutualistic fungi or bacteria to grow (Friesen 2013; Busby et al. 2017), and on animal pollinators for reproduction (Johnson et al. 2015); and the gut microbiome is increasingly recognized as a key aspect of human physiology (Sachs et al. 2011; Shapira 2016). One common feature of most mutualisms is their abundant genetic variation in partner quality—the fitness benefits provided by one partner to another—despite the fact that natural selection is expected to erode variation in mutualism strategies over time (Heath and Stinchcombe 2014). Here we show that partner quality variation in the mutualism between plants and nitrogen-fixing bacteria may be maintained by a genetic tradeoff between attracting mutualistic bacteria and repelling parasitic nematodes.

The maintenance of genetic variation for partner quality in mutualisms is an evolutionary paradox (Heath and Stinchcombe 2014). As with other fitness-related traits, natural selection is expected to drive the highest-fitness partner strategy to fixation, eliminating low-fitness genotypes. Yet genetic variation in partner quality is ubiquitous (Smith and Goodman 1999; Ness et al. 2006; Heath 2010; Hoeksema 2010). Several hypotheses have been advanced to explain this pattern in mutualisms (Heath and Stinchcombe 2014), including the context dependence of partner quality (Barrett et al. 2012; Heath et al. 2012; Simonsen and Stinchcombe 2014a; Burghardt et al. 2017; Harrison et al. 2017b) and frequency-dependent selection balancing cooperative and uncooperative mutualist genotypes (Porter and Simms 2014; Jones et al. 2015). By contrast, the alternative hypothesis that selection favors poor-quality mutualists over high-quality mutualists under some ecological conditions (Bronstein 2001a, b) remains relatively understudied.

Parasites are one agent of selection with the potential to reverse selection on cooperative traits in mutualisms. Parasites can induce major changes in the function and benefits of mutualism, generally in two ways (Strauss and Irwin 2004). First, parasites disrupt the occurrence (i.e., change the frequency) of mutualistic partnerships, typically causing infected hosts to form fewer mutualistic associations (Strauss et al. 2002; De Román et al. 2011; Ballhorn et al. 2014). Second, if the same trait attracts both mutualists and parasites—for example, flowers that draw herbivores to plants along with pollinators—individuals experience a tradeoff between the benefits of mutualism and the costs of parasitism (Gomez 2003; Irwin et al. 2004; Siepielski and Benkman 2009; Ågren et al. 2013; Knauer and Schiestl 2017; Zust and Agrawal 2017). Coupled with spatial or temporal variation in parasite abundance, conflicting selection imposed by mutualists and parasites has been shown to maintain phenotypic variation in mutualism traits (Siepielski and Benkman 2009; Ågren et al. 2013).

We lack direct evidence, however, that tradeoffs between mutualism and parasitism are genetically based, a necessary criterion for selection imposed by parasites to contribute to the maintenance of genetic variation in mutualism (Strauss and Irwin 2004; Heath and Stinchcombe 2014). Genetic trade-offs between mutualism and parasitism can preserve genetic variation in mutualist quality in at least two complementary ways. First, if the genotypes that form the most mutualistic associations (or provide the greatest benefit to their partners) necessarily suffer more parasitism, this may reduce or eliminate their fitness advantage, preventing or slowing the fixation of the ‘best’ mutualist genotypes in populations. Second, if mutualism and parasitism are genetically linked, correlated evolutionary responses may lead to temporally variable selection on mutualism- and parasitism-related traits. That is, if selection favoring effective mutualists causes a correlated decrease in parasite resistance, eventually countervailing selection favoring increased parasite resistance is likely to drive a correlated decrease in mutualist quality, thus preserving variation in mutualism traits. In similar fashion, spatial variation in the abundance of mutualists or parasites can create a mosaic of correlated responses to selection in mutualism- or parasitism-related traits, preserving genetic variation at larger spatial scales. A genetic relationship between mutualism and parasitism traits is one precondition for these evolutionary forces contribute to the maintenance of genetic variation for partner quality in mutualisms.

Although we lack direct evidence for genetic tradeoffs between mutualism and parasitism in most systems, several lines of indirect evidence raise the intriguing possibility that susceptibility to parasites is a common pleiotropic genetic cost of mutualism. Many species are attacked by parasites that bear remarkable resemblance to their mutualists (Adams et al. 2012; Chomicki et al. 2015), and parasites and mutualists frequently use the same cues to infiltrate their host (Sachs et al. 2011). Host genes that affect interactions with mutualists are often also used in defense against parasites (Sachs et al. 2011; Damiani et al. 2012). Consistent with this observation, some species suppress immune function when establishing mutualistic partnerships, leaving them vulnerable to infection (Toth et al. 1990; Miller 1993; Salem et al. 2015). Ultimately, it remains unclear whether these mechanistic tradeoffs create genetic conflict between mutualism and parasitism at the population level, and whether there is genetic variation for the extent to which parasites influence mutualism structure and function.

The keystone ecological and agricultural mutualism between leguminous plants and nitrogen-fixing bacteria (rhizobia) is a promising system for testing for genetic tradeoffs between mutualism and parasitism. In this mutualism, rhizobia provide their plant host with nitrogen, and the plant trades carbohydrates in return. Plants house rhizobia in root organs called nodules (Figure 1). However, many legumes are also infected by parasitic root-knot nematodes that steal photosynthates (Dhandaydham et al. 2008; Goverse and Smant 2014). Nematodes form galls on plant roots that are strikingly similar to the nodules formed by rhizobia (Figure 1). Molecular genetic evidence suggests that genetic conflict between legume responses to mutualistic rhizobia and parasitic nematodes is extensive. Nematodes infiltrate the plant via a stereotyped infection process that mimics that of rhizobia (Goverse and Smant 2014). Many of the same legume genes mediate the two interactions (e.g., receptor genes required to initiate infection) (Koltai et al. 2001; Weerasinghe et al. 2005; Dhandaydham et al. 2008; Damiani et al. 2012). Finally, nematodes have acquired several parasitism genes via horizontal gene transfer from close relatives of rhizobia (Danchin et al. 2010, 2016).

**Figure 1.**
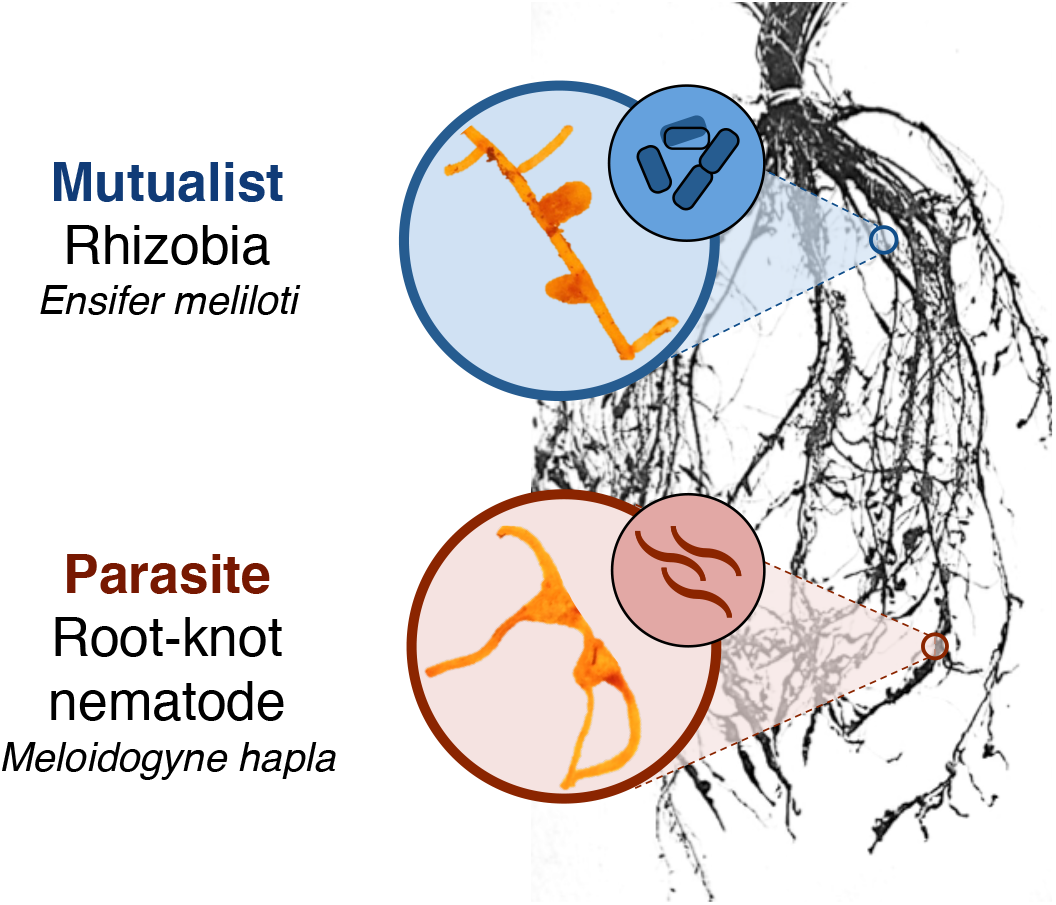
Nodules formed by mutualistic rhizobia (top) and galls formed by parasitic nematodes (bottom) on legume roots. Each gall contains one female nematode. Root image adapted from an image by L.T. Leonard (Fred et al. 1932).

In this report we describe a genetic conflict between plant responses to mutualistic rhizobia and parasitic nematodes in the model legume *Medicago truncatula*. Using two greenhouse-based inoculation experiments, we addressed four questions: (1) How do rhizobia and nematodes impact fitness in co-infected plants?; (2) Is there genetic variation in plant susceptibility to nematodes?; (3) Is there genetic conflict between plant responses to mutualistic rhizobia and parasitic nematodes?; and (4) How do parasitic nematodes impact the rhizobia mutualism? We found a genetic tradeoff between attracting rhizobia and repelling nematodes, and that nematodes disrupt the legume-rhizobia mutualism by deterring nodulation. Our results suggest that genetic conflict with nematodes may maintain variation in partner quality in the legume-rhizobia mutualism, and that genetic tradeoffs with parasitism may be an important overlooked source of variation in positive species interactions.

## Methods

### Study species

*Medicago truncatula* is an annual plant native to the Mediterranean (Cook 1999). Because nodule number is correlated with rhizobia fitness in *M. truncatula* (Heath and Tiffin 2009), it can be used as a proxy for *M. truncatula*'s partner quality (i.e., the benefits it provides) in the rhizobia mutualism. The *M. truncatula* accessions used for this study came from the French National Institute for Agricultural Research (INRA), and the US National Plant Germplasm System (NPGS) Western Regional Plant Introduction Station. For rhizobia inoculations, we used the *E. meliloti* strain Em1022, a highly effective nitrogen-fixer, supplied by (Batstone et al. 2016). We obtained soil infected with the northern root-knot nematode *Meloidogyne hapla* from Dr. Benjamin Mimee (Agriculture and Agri-food Canada, Saint-Jean-sur-Richelieu, Quebec), and maintained these nematodes on tomato plants (cv. Rutgers) in growth chambers and greenhouses at the University of Toronto.

### Greenhouse experiments

We performed two greenhouse experiments to investigate genetic conflict between *M. truncatula*'s response to mutualistic rhizobia and parasitic nematodes. In both experiments, we scarified *M. truncatula* seeds with a razor blade, sterilized them in bleach and ethanol, and stratified them in the dark at 4°C for 36 hours on sterile water agar plates (Garcia et al. 2006). We incubated seeds at room temperature for 16 hours before planting to initiate radicle elongation. We planted the seedlings into sand in 120ml autoclavable Cone-tainers, autoclaved twice at 121°C, and maintained seedlings in the greenhouse at the University of Toronto at 22ºC during the day and 18ºC at night, on a 16:8 light:dark cycle. We top-watered seedlings with distilled water for two weeks, and bottom-watered them for the remainder of the experiments.

Two weeks after germination, we inoculated seedlings with the rhizobium *E. meliloti* and the nematode *M. hapla*. We cultured rhizobia strain Em1022 on solid tryptone yeast (TY) agar media, re-plated colonies three times, and inoculated liquid TY media with these cultures. We diluted liquid cultures to an OD600 reading of 0.1, following previous methods (Simonsen and Stinchcombe 2014b), and inoculated each plant with 1mL of culture at two and three weeks post-germination. We inoculated plants with nematode eggs at the same time. To harvest nematode eggs from infected tomato plants for inoculation, we followed a bleach extraction protocol (Eisenback 2000). Female nematodes lay several hundred eggs into a gelatinous matrix on the outside of each gall (Maggenti and Allen 1960). We rinsed the roots of infected tomato plants and incubated them in a shaker at room temperature for 5 minutes in 10% commercial bleach (0.5% NaOCl) to dissolve the gelatinous matrix binding the eggs together. We poured the solution through a series of mesh soil sieves, collected nematode eggs on a #500 mesh sieve (25µm pore size), and stored collected eggs in distilled water in Falcon tubes. We inoculated each plant with nematode eggs twice (at two and three weeks post-germination), on the same schedule as the rhizobia inoculations.

*Experiment 1*: To test how rhizobia and nematodes impact fitness in co-infected plants, and to measure genetic variation in nematode infectivity, we used a fractional factorial design with a total of 400 *M. truncatula* plants from 10 genotypes across 10 blocks. We inoculated each plant with 1 of 40 nematode genotypes. Each block included 4 replicates of each plant genotype and 1 replicate of each nematode genotype, for a total of 40 replicates of each plant genotype and 10 replicates of each nematode genotype. Each nematode genotype inoculated 2 different plant genotypes, for a total of 5 replicates per nematode genotype-plant genotype combination.

To culture individual nematode genotypes, we inoculated tomatoes with second-stage juvenile (J2) nematodes from individual egg masses (Thies et al. 2002). This protocol ensured that each tomato plant was infected by a single maternal family. After approximately three months, we extracted nematode eggs from these tomatoes and used them to inoculate our experimental plants. We inoculated each plant with ~200–400 nematode eggs, depending on availability, and included number of eggs as a covariate in our statistical analyses. Nine plants received >400 eggs; excluding these plants from the analysis did not qualitatively affect our results. We harvested plants 3.5 months after planting.

*Experiment 2*: To measure genetic conflict between attracting rhizobia and repelling nematodes, and to test how parasitic nematodes impact the rhizobia mutualism, we used a split-plot randomized design. Each block contained two treatments: one in which we only inoculated plants with rhizobia, and one in which we inoculated plants with both rhizobia and nematodes. Plants received a total of 400 nematode eggs from a genetically variable nematode inoculum. Each treatment in each block contained one *M. truncatula* individual from each of 50 genotypes. In each block, we bottom-watered all plants in the same treatment from the same tray. We replicated this design across 10 blocks (50 plants per treatment per block × 2 treatments × 10 blocks = 1000 plants). We did not include a nematode-only treatment because *Medicago* grows poorly under nitrogen-poor conditions without rhizobia (Harrison et al. 2017a). We harvested plants 4.5 months after planting.

We checked flowering and collected ripe fruit daily throughout both experiments. Upon harvesting the plants, we stored the roots at 4°C in zip-top plastic bags until processing. We dried the aboveground tissue in a drying oven for approximately 1 week and weighed it to the nearest 1mg. We weighed all fruit each plant produced to measure total fruit mass. To verify that fruit mass was an accurate measurement of reproductive success, we measured the correlation between fruit mass and seed number for a subset of plants (N = 167) and found that fruit mass and seed number were tightly correlated (r = 0.76, P < 0.001, df = 165). We counted the number of nodules and galls on each root system under a dissecting microscope. To capture differences in nodule size, we haphazardly harvested up to ten of the largest nodules on each plant. Nodules were stored in 2mL tubes containing silica desiccant and synthetic polyester for a month until they dried out, and we weighed the dried nodules collected from each plant to the nearest 1µg. We estimated total nodule biomass for each plant by multiplying total nodule number by mean nodule mass. After counting nodules and galls and harvesting nodules, we dried the roots in a drying oven for approximately 1 week and weighed them to the nearest 1mg.

### Statistical analyses

We performed all analyses in R 3.3.2 with deviation coding ("contr.sum") for categorical variables (R Core Team 2016). Unless stated otherwise, we ran all analyses with the *(g)lmer* function in the *lme4* package (Bates et al. 2015). We tested significance of fixed effects with type III sums of squares using the *Anova* function in the *car* package (Fox and Weisberg 2011), and used likelihood ratio tests to test significance of random effects (Bolker et al. 2009). We confirmed that all models met the parametric statistical assumptions of normality, homoscedasticity, and linearity by inspecting quantile-quantile plots, scale-location plots, and plots of the residuals versus fitted values, respectively. We also checked for overdispersion by testing whether the ratio of the residual variance to the residual degrees of freedom was equal to 1. We calculated least-squares treatment and genotype means using the *lsmeans* package (Lenth 2016) and created figures using *ggplot2* (Wickham 2009).

### Effect of rhizobia and nematodes on fitness in co-infected plants (Experiment 1)

To test how rhizobia and nematodes impact fitness in co-infected plants, we analyzed two fitness components, aboveground biomass and total fruit mass. These models included number of nodules, number of galls, root mass, researcher (to control for differences among researchers in nodule and gall counts), and the number of nematode eggs in the inoculum as fixed effects, and block as a random effect. We log-transformed aboveground biomass for analysis. We included a fixed effect of root mass in this and subsequent analyses to control for differences in overall root system size and foraging ability, as well as differences in the root space available for the formation of symbiotic structures (i.e., nodules and galls).

### Genetic variation in plant susceptibility to nematodes (Experiments 1 and 2)

In Experiment 1, we tested for genetic variation in infectivity among nematode genotypes, and for a plant genotype-by-nematode genotype interaction. A genotype-by-genotype interaction for gall number would indicate that the number of galls formed depends on the combination of plant and nematode genotypes. In this analysis, we included random effects of plant genotype, nematode genotype, plant genotype × nematode genotype, and block. We included fixed effects of root mass, researcher (to control for differences among researchers in gall counts), and the number of nematode eggs in the inoculum. We log-transformed gall number for this analysis because the log transformation met parametric statistical assumptions much better than a Poisson or negative binomial GLMM. When testing for the genotype-by-genotype interaction, we excluded plant genotype-nematode genotype combinations with fewer than three replicates.

In Experiment 2, we tested for genetic variation in plant susceptibility to nematodes by testing for significant variation among plant genotypes in the number of galls they produced. This analysis included fixed effects of root mass and researcher (to control for differences among researchers in gall counts), and random effects of plant genotype and block. Gall number was zero-inflated and overdispersed, so we fit a zero-inflated negative binomial GLMM using the R package *glmmADMB* (Fournier et al. 2012).

### Genetic conflict between attracting rhizobia and repelling nematodes (Experiment 2)

To test for genetic conflict between plant responses to mutualistic rhizobia and parasitic nematodes, we estimated the genetic correlation between nodule number and gall number. To estimate genotype means for gall number, we extracted the conditional modes of the genotype random effect from a model that included fixed effects of root mass and researcher, and random effects of genotype and block. Because we found evidence that nematodes disrupt the mutualism by inhibiting nodulation (see Results), we use estimates of nodulation from the rhizobia-only treatment to estimate the genetic correlation with gall formation. We estimated genotype means for nodule number using a model similar to the gall model, and specified a negative binomial error distribution and allowed for zero inflation in both models.

We also estimated the genetic correlation between gall number and the change in nodule number between the two treatments. We estimated genotype means for nodule number in nematode-infected plants using a similar model to the one used to estimate nodule number in the rhizobia-only treatment. We subtracted the genotype mean for nodule number in nematode-infected plants from the genotype mean for nodule number in uninfected plants to calculate the change in nodule number for each genotype.

### Effect of nematodes on the rhizobia mutualism (Experiment 2)

To test how parasitic nematodes impact the rhizobia mutualism, we compared nodule number, mean nodule mass, and total nodule biomass between nematode-infected and uninfected plants. These analyses included treatment (nematode presence or absence) and root mass as fixed effects, and random effects of genotype, block, treatment × genotype and treatment × block. The treatment × block interaction is necessary when analyzing split-plot experiments to allow the effect of nematode treatment to vary across blocks (Altman and Krzywinski 2015). We specified a negative binomial error distribution for nodule number and allowed for zero inflation using the function *glmmadmb* in the R package *glmmADMB* (Fournier et al. 2012), and log-transformed mean nodule mass and total nodule biomass for analysis. The nodule number analysis included a fixed effect of researcher to control for researcher differences in nodule counts.

We ran similar analyses to compare aboveground biomass, flowering time, and total fruit mass between nematode-infected and -uninfected plants. We log-transformed all three variables for analysis, and omitted the fixed effect of root mass. For flowering time and total fruit mass, we analyzed a subset of genotypes (N = 22) with at least three replicates that flowered and fruited in each treatment, to test for treatment × genotype interactions.

## Results

### Effect of rhizobia and nematodes on fitness in co-infected plants (Experiment 1)

Rhizobia and nematodes affected different fitness components in co-infected plants (Figure 2). Plants that formed more nodules had significantly greater aboveground biomass than plants with fewer nodules (χ^2^_df=1_ = 33.918, P < 0.001; Figure 2A). There was no corresponding effect of gall number on aboveground biomass (χ^2^_df=1_ = 0.370, P = 0.543; Figure 2B). By contrast, the number of nodules did not significantly affect the total fruit mass that plants produced (χ^2^_df=1_ = 0.490, P = 0.484; Figure 2C), but plants with more galls produced less total fruit mass than plants with fewer galls (χ^2^_df=1_ = 9.394, P = 0.002; Figure 2D).

**Figure 2.**
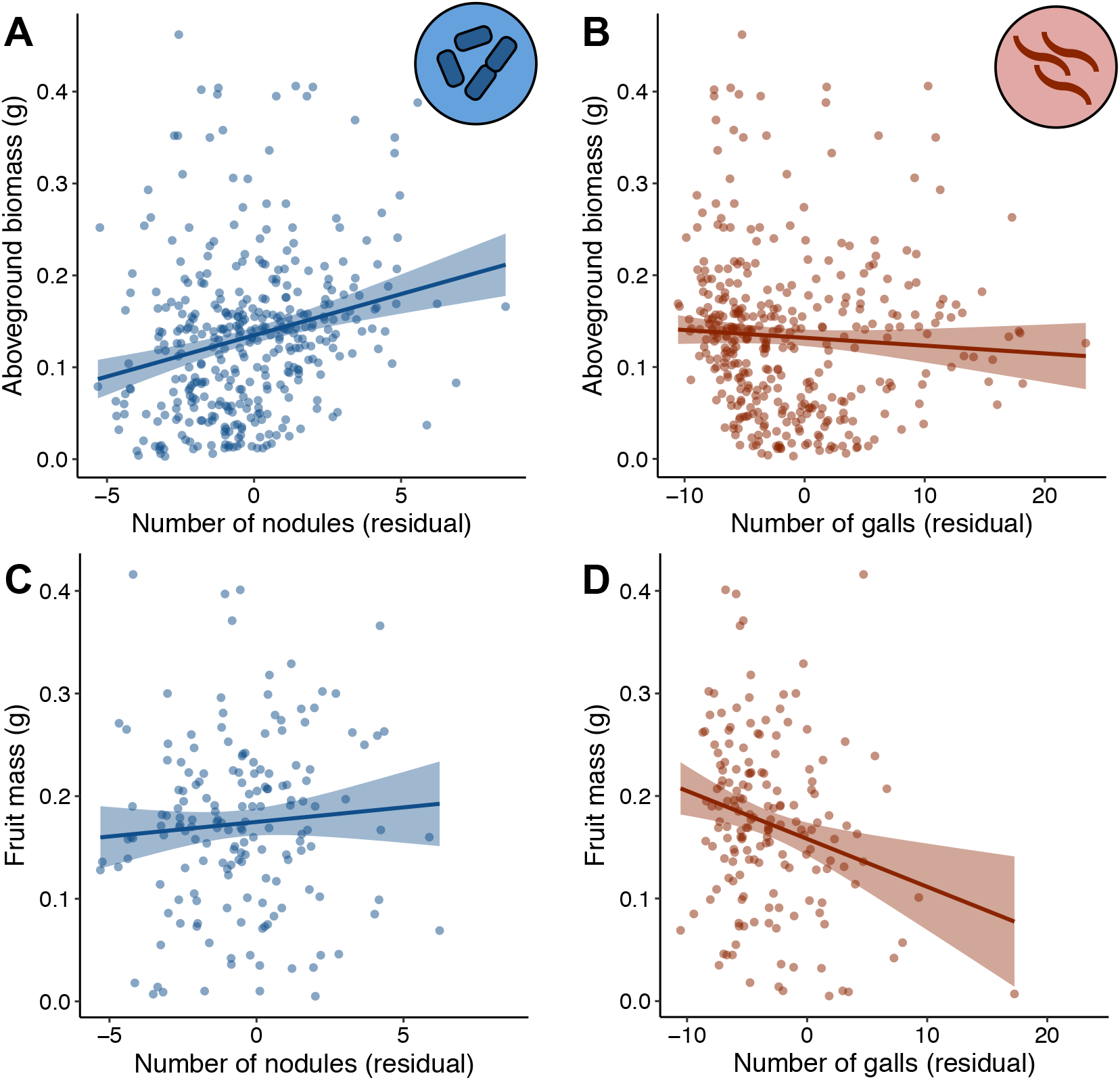
Rhizobia and nematodes affect different plant fitness components in co-infected plants. (A and C) The relationship between nodule number and aboveground biomass (A), and nodule number and fruit mass (C). (B and D) The relationship between gall number aboveground biomass (B), and gall number and fruit mass (D). Bands are standard errors. The negative relationship in (D) remained significant when the point in the lower right-hand corner was removed.

## Genetic variation in plant susceptibility to nematodes (Experiments 1 and 2)

In both experiments, there was significant variation among plant genotypes in the number of galls formed (controlling for root biomass) (Experiment 1: N _genotypes_ = 10, P = 0.001; Experiment 2: N _genotypes_ = 48, P < 0.001), indicating that there is genetic variation in plant susceptibility to nematode infection. In addition, there was significant variation in gall number among nematode genotypes in Experiment 1 (N _genotypes_ = 40, P < 0.001). There was no significant plant genotype × nematode genotype interaction (N _plant-nematode combinations_ = 74, P = 0.539).

### Genetic conflict between attracting rhizobia and repelling nematodes (Experiment 2)

There was a significant positive correlation between gall number and the number of nodules produced in the absence of nematodes (r = 0.30, P = 0.039; Figure 3A). This correlation disappeared when the outlier genotype HM170, which formed 2.9 standard deviations more nodules than the mean in our experiment, was included in the analysis (r = 0.06, P = 0.710). In another study of nodulation in *M. truncatula*, this genotype also formed more nodules than 90% of 250 accessions surveyed (Stanton-Geddes et al. 2013a,b). Together, our results and those of Stanton-Geddes et al. suggest that this genotype may be a biological outlier with respect to the rhizobia mutualism, so we ran subsequent analyses with and without this outlier genotype.

**Figure 3.**
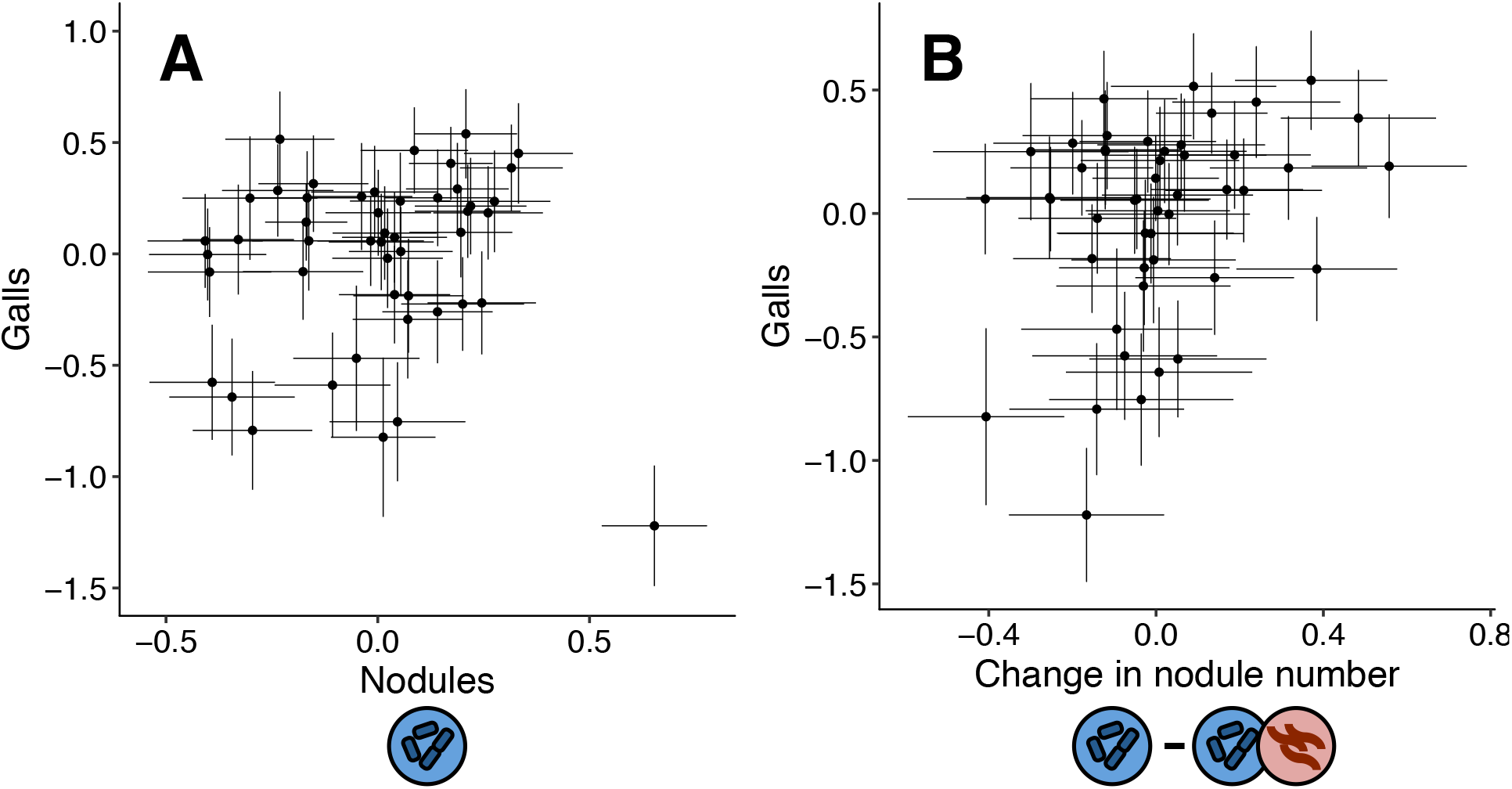
Genetic correlation between the number of galls and nodules that plants produce. Points are conditional modes (BLUPs) for each plant genotype ± SE. (A) Genetic correlation between gall number and the number of nodules produced by plants in the absence of nematodes. There is a significant positive correlation when the outlier in the lower right-hand corner is excluded (r = 0.30, P = 0.039), but not when it is included (r = 0.06, P = 0.710). (B) Genetic correlation between gall number and the change in nodule number in the absence and presence of nematodes (r = 0.31, P = 0.034). Excluding HM170 did not qualitatively change this result (r = 0.29, P = 0.052). We used a resampling approach to generate the standard errors on the change in nodule number in panel B.

There was no significant genetic correlation between gall number and the number of nodules produced in the presence of nematodes, regardless of whether the outlier genotype HM170 was included in the analysis (with HM170: r = −0.20, P = 0.153; without HM170: r = 0.04, P = 0.789). However, there was a significant positive genetic correlation between gall number and the change in nodule number between the two treatments (r = 0.31, P = 0.034; Figure 3B), indicating that plant genotypes that were most susceptible to nematodes (i.e., formed the most galls) decreased the most in nodule number when infected with nematodes. Excluding HM170 did not qualitatively change this result (r = 0.29, P = 0.052).

### Effect of nematodes on the rhizobia mutualism (Experiment 2)

Nematode-infected plants produced fewer nodules and less total nodule biomass than uninfected plants, although mean nodule mass did not differ between infected and uninfected plants (Table 1, Figure 4A-C). There was a significant effect of plant genotype for all nodule traits (Table 1), indicating that genotypes differed in mutualism phenotypes. We detected a significant treatment × genotype interaction for nodule number and a marginally significant treatment × genotype interaction for total nodule biomass (Table 1, Figures 4D & 4F). These interactions indicate that plant genotypes differed in how nodule traits were impacted by nematode infection. There was no treatment × genotype interaction for mean nodule mass (Table 1, Figure 4E). Our results were qualitatively similar when we removed the outlier genotype HM170 (see Figure 4D).

**Table 1.**
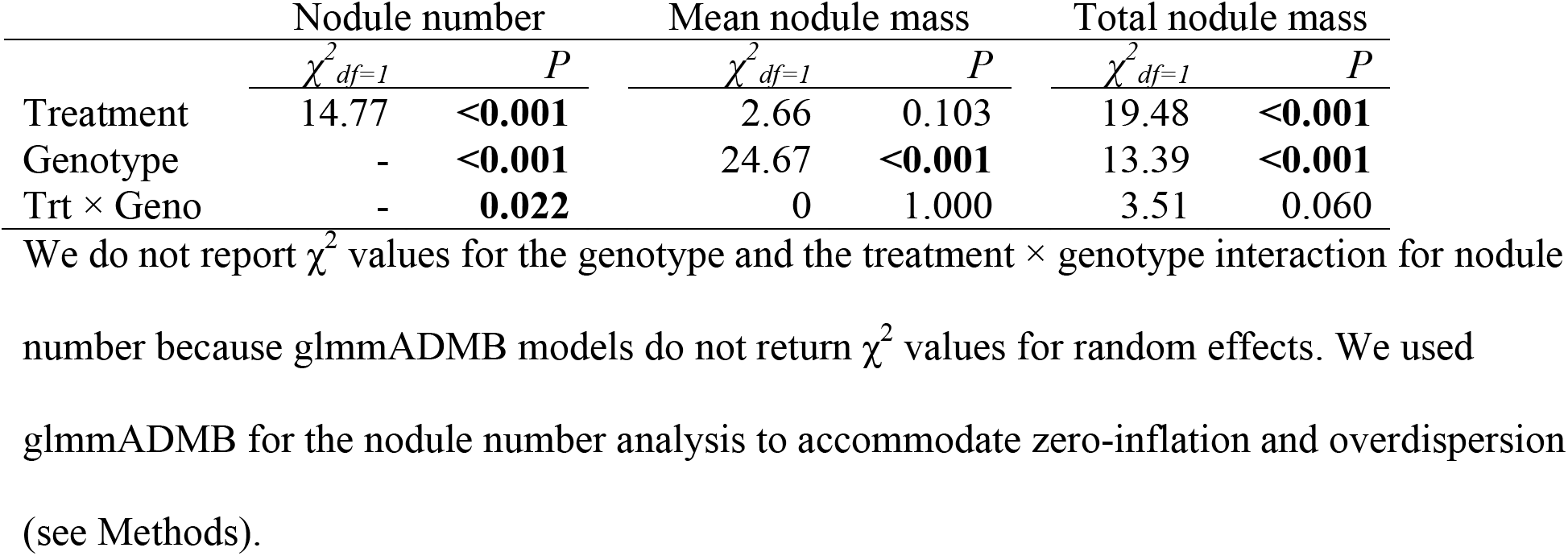
Effect of treatment (nematode presence or absence), plant genotype, and the treatment × genotype interaction on nodule traits.

|  | Nodule number |  | Mean nodule mass |  | Total nodule mass |  |
| --- | --- | --- | --- | --- | --- | --- |
| | $\chi^2_{df=1}$ | *P* | $\chi^2_{df=1}$ | *P* | $\chi^2_{df=1}$ | *P* |
| Treatment | 14.77 | **<0.001** | 2.66 | 0.103 | 19.48 | **<0.001** |
| Genotype | - | **<0.001** | 24.67 | **<0.001** | 13.39 | **<0.001** |
| Trt × Geno | - | **0.022** | 0 | 1.000 | 3.51 | 0.060 |

We do not report $\chi^2$ values for the genotype and the treatment × genotype interaction for nodule number because glmmADMB models do not return $\chi^2$ values for random effects. We used glmmADMB for the nodule number analysis to accommodate zero-inflation and overdispersion (see Methods).

**Figure 4.**
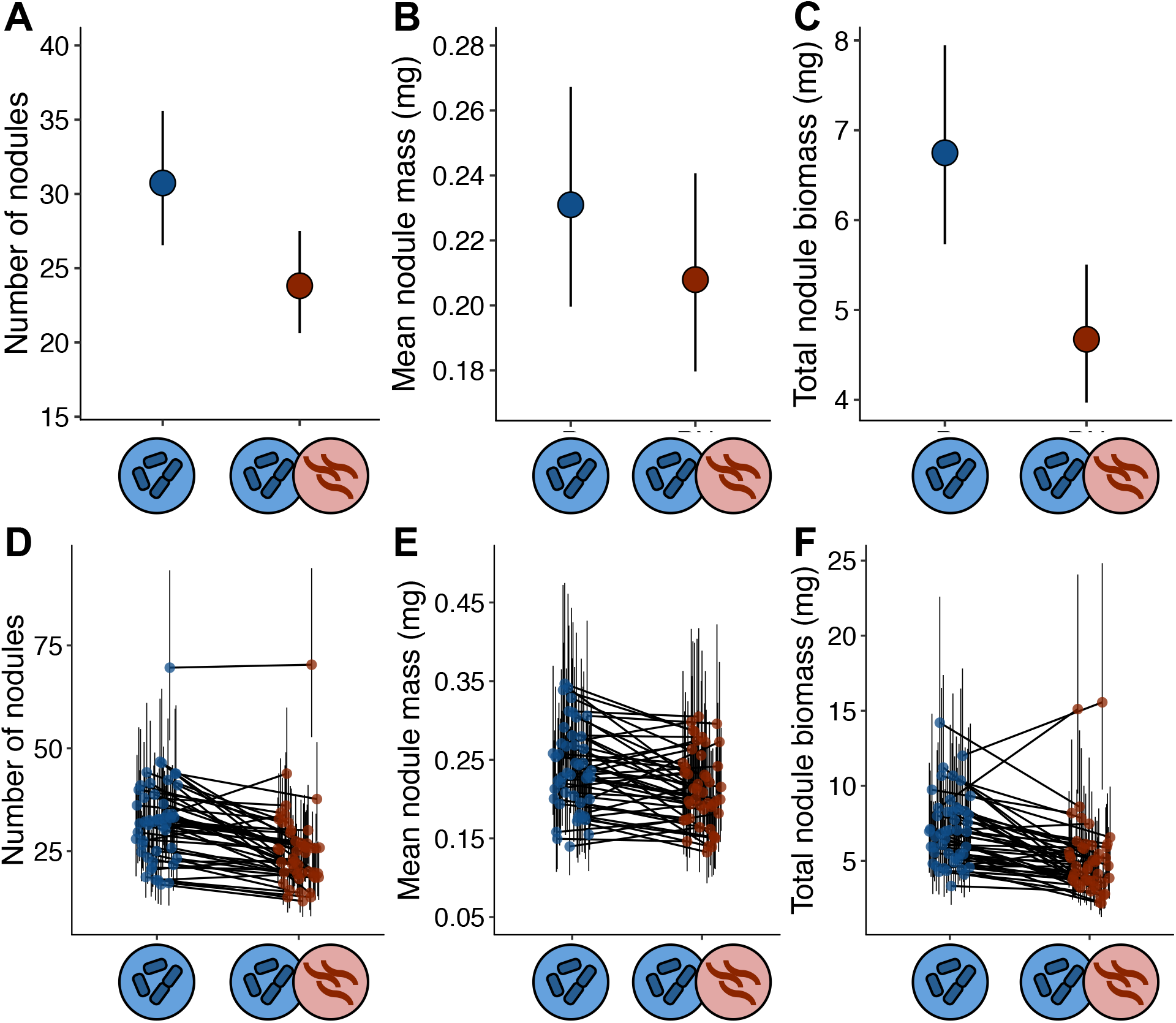
Nematodes affect the nodule phenotypes. Number of nodules (A), mean nodule mass (B), and total nodule biomass (number of nodules × mean nodule mass) in nematode-infected and -uninfected plants. In A-C, points are least-squares treatment means ±95% CIs. (D-F) Genotype-by treatment interactions for number of nodules (D), mean nodule mass (E), and total nodule biomass (F). In each treatment, points are least-squares genotype means ±95% CIs; lines connect the same genotype in the two treatments.

Other plant traits were not strongly affected by nematode infection. Although there was significant genetic variation for all plant traits, there was no difference between infected and uninfected plants in aboveground biomass, flowering time, or total fruit mass (Table 2). There was a marginally significant treatment × genotype interaction for aboveground biomass (Table 2).

**Table 2.** Effect of treatment (nematode presence or absence), plant genotype, and the treatment × genotype interaction on plant traits. Top: Aboveground biomass, flowering time, and total fruit mass.

|  | Aboveground biomass |  | Flowering time |  | Total fruit mass |  |
| --- | --- | --- | --- | --- | --- | --- |
| | $\chi^2_{df=1}$ | *P* | $\chi^2_{df=1}$ | *P* | $\chi^2_{df=1}$ | *P* |
| Treatment | 0.47 | 0.495 | 0.66 | 0.417 | 2.76 | 0.096 |
| Genotype | 56.33 | **<0.001** | 17.90 | **<0.001** | 5.65 | **<0.001** |
| Trt × Geno | 3.55 | 0.059 | 0 | 1.000 | 0.03 | 0.859 |

## Discussion

Here we showed that an ecologically relevant parasite disrupts the mutualism between leguminous plants and nitrogen-fixing rhizobial bacteria. *Medicago truncatula* plants that were infected by parasitic nematodes formed fewer rhizobia nodules and less nodule biomass per gram of root tissue than uninfected plants. Strikingly, nematode infection impacted nodule traits more strongly than other plant phenotypes, indicating that the parasite's effect on the legume-rhizobia mutualism is not merely a byproduct of lower overall performance in infected plants. Moreover, we found that a plant's affinity for rhizobia and susceptibility to nematodes were genetically correlated: plants that formed more nodules with rhizobia were more heavily infected by nematodes. Our results suggest that genetic conflict with parasitic nematodes is an important factor shaping the *Medicago*-rhizobia mutualism. If genetic conflict with parasitism is a general feature of many mutualisms, it may contribute to the maintenance of genetic variation for partner quality and influence evolution in positive species interactions.

### *Nematodes decrease associations between* Medicago *and mutualistic rhizobia*

Our work extends past research on the impact of antagonists on mutualism in two key ways. First, we showed that mutualism traits were more strongly impacted by parasite infection than other aspects of plant performance. In the presence of nematodes, plants formed fewer associations with mutualistic rhizobia. We found that nematode-infected *Medicago* plants formed 23% fewer nodules and 19% less total nodule biomass per gram of root than uninfected plants (Figure 4A-C). By contrast, nematode infection only weakly affected aboveground biomass, flowering time, and total fruit mass (Tables 1 and 2). Future work in other mutualisms should explore whether elevated sensitivity to parasites is a characteristic feature of mutualism traits.

Second, when ecological factors influence mutualistic associations, their evolutionary consequences depend on whether there is standing genetic variation for environmental responsiveness in the form of genotype-by-environment interactions. Although environmental effects on mutualism are common (Bronstein 1994; Bronstein et al. 2003; Kersch and Fonseca 2005; Afkhami et al. 2014), and antagonists often interfere with the fitness benefits of mutualists (Gomez 2005; Liere and Larsen 2010; Simonsen and Stinchcombe 2014a), these effects are rarely investigated from a genetic perspective (but see (Heath et al. 2010)).

Our results demonstrate that there is standing genetic variation for *Medicago*'s susceptibility to parasite infection, as well as in the degree to which the plant's mutualism was robust to parasite-mediated disruption (treatment × genotype interaction: Table 1 and Figure 4D-F). *Medicago truncatula* genotypes varied significantly in their susceptibility to nematode infection, with some genotypes forming dozens or hundreds of galls while others formed few or none. Moreover, while some plant genotypes formed substantially fewer nodules when infected by nematodes, others—including one hyper-nodulating outlier (Figure 4D)—were largely unaffected by the parasite. The degree to which the *Medicago*-rhizobia mutualism is impacted by parasitic nematodes therefore has the genetic capacity to evolve. There was also genetic variation in infectivity in the nematode population (i.e., nematode genotypes differed in the number of galls they formed on plant roots), demonstrating that both the plant and the parasite have the genetic capacity to evolve in response to the other. However, we found no evidence for genotype-by-genotype interactions between plants and nematodes that would facilitate coevolution in the system.

### Genetic tradeoff between attracting a mutualist and repelling a parasite

*Medicago truncatula*'s susceptibility to nematode infection was genetically correlated with its affinity for mutualistic rhizobia (Figure 3A). Plant genotypes that formed the most rhizobia nodules also formed the most galls, while genotypes that formed few nodules were more resistant to nematode infection. One caveat to this result is that the genetic correlation was no longer significant when the hyper-nodulating outlier genotype was included (Figure 3A). This outlier appears to be behaving fundamentally differently with respect to the rhizobia mutualism, and may be an informative point of comparison for future work on the genomic underpinnings of the genetic correlation between nodulation and galling.

The genetic correlation between attracting rhizobia and repelling nematodes in *Medicago* is consistent with molecular genetic work in the legume *Lotus japonicus* showing that mutants that do not form nodules are also resistant to nematode infection (Weerasinghe et al. 2005). To our knowledge, only a handful of past studies have documented genetic conflict between mutualism and parasitism (Toth et al. 1990; Miller 1993). Both examined the symbiosis between plants and mycorrhizal fungi, and found pathogen-resistant genotypes formed fewer mycorrhizal associations. Genetic conflict may be a general feature of intimate symbioses like plant-microbe mutualisms, in which one partner lives inside the tissue of another.

A genetic correlation underlying the tradeoff between attracting mutualists and repelling parasites, like the one we documented in *M. truncatula,* is one mechanism that can contribute to the maintenance of genetic variation for partner quality in mutualisms (Heath and Stinchcombe 2014) and alter evolutionary trajectories (Nuismer and Doebeli 2004; Strauss and Irwin 2004). The genetic tradeoff between mutualism and parasitism has distinct evolutionary consequences for mutualism at different spatial and temporal scales. First, within a single population, as more mutualistic genotypes spread, susceptibility to parasites is also spreading: eventually, this should erode, or eliminate the fitness advantage gained by being a better mutualist partner, slowing or preventing their fixation. Second, variation in mutualist and parasite abundance among sites or years is likely to create a selection mosaic that favors high-quality partners where parasites are absent and low-quality partners where parasites are absent. Such spatial and temporal variation in the direction of selection and could maintain genetic variation for partner quality in mutualism (Thompson 2005; Huang et al. 2015). To directly assess how the genetic tradeoffs we report influences selection on, and variation in, partner quality in the legume-rhizobia mutualism, future work should characterize spatial and temporal variation in nematode and rhizobia abundance in wild *Medicago* populations.

Although there are surprisingly few estimates of the genetic tradeoff between mutualism and antagonism, widespread tradeoffs at the phenotypic level suggest that genetic conflict between positive and negative species interactions may be extremely common. For example, trypanosomatid parasites of firebugs mimic the vertical transmission mechanisms of their host's bacterial symbionts, such that symbiont transmission is associated with a risk of parasite infection (Salem et al. 2015). In the seed dispersal mutualism between Clark's nutcracker and pine trees, selection exerted by a seed predator opposes mutualist-mediated selection (Siepielski and Benkman 2009). Pollinators and herbivores often cue in on the same plant signals, imposing conflicting selection on floral displays that weakens the overall strength of pollinator-mediated selection (Rey et al. 2002; Gomez 2003; Schiestl et al. 2011, 2014; Ågren et al. 2013; Knauer and Schiestl 2017). If these phenotypic tradeoffs are underpinned by genetic correlations like the correlation we report in the legume-rhizobia mutualism, genetic conflict with parasitic interactions is likely an important source of genetic variation in diverse mutualistic systems (Bronstein 2001a; Strauss and Irwin 2004).

### Implications for mutualism evolution

Our study joins a number of others demonstrating that an evolutionary genetic approach to mutualism can yield meaningful new insights about these positive species interactions (Heath 2010; Heath et al. 2012; Afkhami and Stinchcombe 2016; Burghardt et al. 2017). A recent transcriptomic study of the legume-rhizobia mutualism, for example, showed that genotype-by-genotype interactions between plants and rhizobia impact carbon and nitrogen exchange, the central function of the symbiosis (Burghardt et al. 2017). Intriguingly, Burghardt et al. (2017) also found significant variation among plant genotypes in the expression of defense genes in nodules. Together, our study and theirs raise the possibility that conflict with plant immunity is a key feature of the legume-rhizobia mutualism whose evolutionary significance has been largely overlooked.

Genetic conflict with parasites could significantly alter the rate and trajectory of evolution in mutualisms. The impact of this conflict on mutualism evolution depends on three factors about which little is known in any system: the degree of overlap in the genetic pathways controlling the two symbioses; how parasites disrupt mutualistic partnerships; and the ecological factors that mediate conflict. All three of these factors warrant further study in the legume-rhizobia mutualism, and a diverse array of other positive species interactions.

## Acknowledgements

We are grateful to Benjamin Mimee at Agriculture and Agri-food Canada for providing nematode inoculum, and Dennis VanDyk at the University of Guelph's Muck Crops Research Station for providing tomato cv. Rutgers seeds and advice on nematode culturing. Rebecca Batstone provided the rhizobia strain Em1022 used in these experiments, as well as advice on experimental protocols. Thanks to Bill Cole, Andrew Petrie, and Bruce Hall for greenhouse assistance, and Amalia Caballero, Monina Cepeda, Lily He, and Diana Nakib for help with data collection. This work was funded by an NSERC grant to John Stinchcombe and a University of Toronto EEB Postdoctoral Fellowship to Corlett Wood.

## Author contributions

CW and JS conceived of the study, and CW, BP, PV and JS designed the experiments. CW, BP, PV, and CB performed the experiments and collected data. All authors contributed to data analysis and interpretation. CW wrote the initial draft of the manuscript, and all authors contributed to manuscript revisions.

## Data accessibility

Data from both experiments will be available on Dryad.

